# Chromosome size-dependent polar ejection force impairs mammalian mitotic error correction

**DOI:** 10.1101/2023.10.16.562637

**Authors:** Megan K. Chong, Miquel Rosas-Salvans, Vanna Tran, Sophie Dumont

## Abstract

Accurate chromosome segregation requires sister kinetochores to biorient, attaching to opposite spindle poles. To this end, the mammalian kinetochore destabilizes incorrect attachments and stabilizes correct ones, but how it discriminates between these is not yet clear. Here, we test the model that kinetochore tension is the stabilizing cue and ask how chromosome size impacts that model. We live image PtK2 cells, with just 14 chromosomes, widely ranging in size, and find that long chromosomes align at the metaphase plate later than short chromosomes. Enriching for errors and imaging error correction live, we show that long chromosomes exhibit a specific delay in correcting attachments. Using chromokinesin overexpression and laser ablation to perturb polar ejection forces, we find that chromosome size and force on arms determine alignment order. Thus, we propose a model where increased force on long chromosomes can falsely stabilize incorrect attachments, delaying their biorientation. As such, long chromosomes may require compensatory mechanisms for correcting errors to avoid chromosomal instability.

## Introduction

The kinetochore is the multivalent interface that connects chromosomes to spindle microtubules at cell division. Accurate chromosome segregation requires biorientation, a state in which sister kinetochores attach to opposite spindle poles. To preserve genome integrity, each kinetochore must monitor chromosomes’ attachment status and signal to the cell whether or not it is ready to enter anaphase (Musacchio and Desai, 2017). Correct, bioriented attachments form alongside incorrect attachments, which must be detected and corrected through a process called error correction. Both physical and biochemical features differ between correct and incorrect attachments and understanding how these cues govern error correction and how their detection varies across chromosomes is central to understanding mitotic fidelity (Lampson and Grishchuk, 2017).

Current models for error correction propose that incorrect kinetochore-microtubule attachments are molecularly destabilized to promote detachment, while correct attachments are stabilized (Funabiki, 2019; Sarangapani and Asbury, 2014). A key regulator in this process is Aurora B kinase, which phosphorylates the kinetochore’s primary loadbearing complex Ndc80C (Powers et al., 2009) to reduce its affinity for microtubules (Biggins and Murray, 2001; Cheeseman et al., 2006; DeLuca et al., 2011; Zaytsev et al., 2015). This phosphorylation decreases with kinetochore tension and is required for error correction to occur (DeLuca et al., 2006; Lampson et al., 2004). Moreover, applying ectopic tension to incorrect attachments is sufficient to delay their correction indefinitely in grasshopper spermatocytes (Nicklas and Koch, 1969), indicating a causative role for tension in distinguishing between correct and incorrect attachments in some cell types. However, how the mammalian error correction machinery responds to acute changes in tension and what cellular sources of said tension are relevant remain poorly understood.

While tension at bioriented kinetochores is generated primarily by kinetochore-bound microtubules (k-fibers) exerting opposing poleward pulling force on sisters, non-kinetochore microtubules also exert force in the spindle that influences kinetochore behavior. Polar ejection force arises from non-kinetochore microtubules growing outward from spindle poles generating outward force on chromosome arms (Rieder et al., 1986). This polar ejection force, mediated both by microtubule polymerization and by two chromokinesins in mammalian cells (Kif22/Kid and Kif4a), contributes to chromosome congression and regulates the amplitude of kinetochore oscillations at metaphase (Barisic et al., 2014; Iemura and Tanaka, 2015; Ke et al., 2009; Levesque and Compton, 2001; Wandke et al., 2012). Overexpression of the Drosophila orthologue of Kif22/Kid induces attachment errors, which elude error correction (Cane et al., 2013b), suggesting that polar ejection forces can contribute to attachment stabilization, possibly by increasing kinetochore tension. In mammalian cells, tension perturbations are needed to understand how the error correction machinery responds to tension changes and how the magnitude of polar ejection force—which in principle can vary throughout mitosis (Ke et al., 2009; Thompson et al., 2022) and across different chromosomes—impacts kinetochore tension and attachment stability.

Segregation error rates vary not only across cell types but also across chromosomes within the same cell type. Indeed, long chromosomes have been shown to missegregate more frequently than short chromosomes in human cells (Klaasen et al., 2022; Worrall et al., 2018). Many mechanisms could underlie this difference including differential nuclear positioning in interphase (Klaasen et al., 2022), distinct positions of long and short chromosomes along the metaphase plate (Rieder and Salmon, 1994; Wan et al., 2012), or variable biophysical features between long and short chromosomes. Indeed, given the biophysical nature of error detection models, error correction cues might be read differently based on the physical features of the chromosome or kinetochore in question, complicating our understanding of how tension impacts kinetochore-microtubule attachment stability. However, studying segregation outcomes is not sufficient to understand the dynamic formation of correct attachments. Instead, probing how efficiently chromosomes of different sizes become correctly bioriented sheds light not only on the cues driving error correction, but also on the mechanisms underlying chromosome-based differences in mitotic trajectories and segregation outcomes.

Here, we provide direct evidence for the tension model for error correction in mammalian cells and show that differential tension at chromosomes of different sizes impacts both their biorientation and error correction efficiency. In mammalian rat kangaroo (PtK2) cells, we show that long chromosomes become bioriented later and experience higher pushing force than short chromosomes in the same cell. By enriching for errors and letting them correct in live imaging of drug washouts, we show that long chromosomes are less efficient at correcting errors therefore delaying their biorientation. Increasing size-based force via chromokinesin Kif22/Kid overexpression increases the probability of chromosomes of any size becoming persistently stuck at spindle poles. Finally, with laser ablation, we cut chromosome arms and find that chromosome size, not identity, sets biorientation efficiency. We propose a model in which elevated force on long chromosomes increases kinetochore tension, stabilizing kinetochore-microtubule attachments, thus prolonging the lifetime of erroneous attachments at long chromosomes. These findings provide a framework for understanding not just mechanisms of error detection, which may vary in response to chromosome-specific differences, but also aneuploidy and karyotype evolution.

## Results and Discussion

### Long chromosomes align less efficiently than short chromosomes and experience higher spindle pushing force

To determine which physical features promote correct kinetochore-microtubule attachment in mitosis, and whether they vary across chromosomes, we assessed whether dividing rat kangaroo (PtK2) cells exhibit a bias in chromosome alignment efficiency. With just 14 chromosomes, widely ranging in size, PtK2 cells are particularly well suited to this question. We classified chromosomes by size into two, roughly even groups for easy identification: long chromosomes (≥7μm along the longest dimension by phase microscopy) and short ones (<7μm) (Fig. 1A & 1B). We imaged live spindle assembly in PtK2 cells expressing eYFP-Cdc20 to mark kinetochores and scored the frequency of early and late aligning chromosomes belonging to either the long or short chromosome group. Here, we define early aligning chromosomes as the first three chromosomes to begin oscillating at the metaphase plate, a marker of biorientation, and late aligning chromosomes as the last three chromosomes to align (Fig. 1C). While long and short chromosomes were evenly represented in the early aligning group, long chromosomes were significantly overrepresented among late aligning chromosomes (Fig. 1D, 1E, and Video 1; Fisher’s exact test p=0.0041). Thus, on average, long chromosomes biorient and form correct attachments later in spindle assembly than short chromosomes in the same cell, suggesting less efficient or delayed attachment formation. Such a delay could be due to chromosome position differences in the spindle, intrinsic differences in chromosome or kinetochore identity, and/or differences in the formation or correction of incorrect attachments that must be resolved prior to alignment.

**Figure 1.**
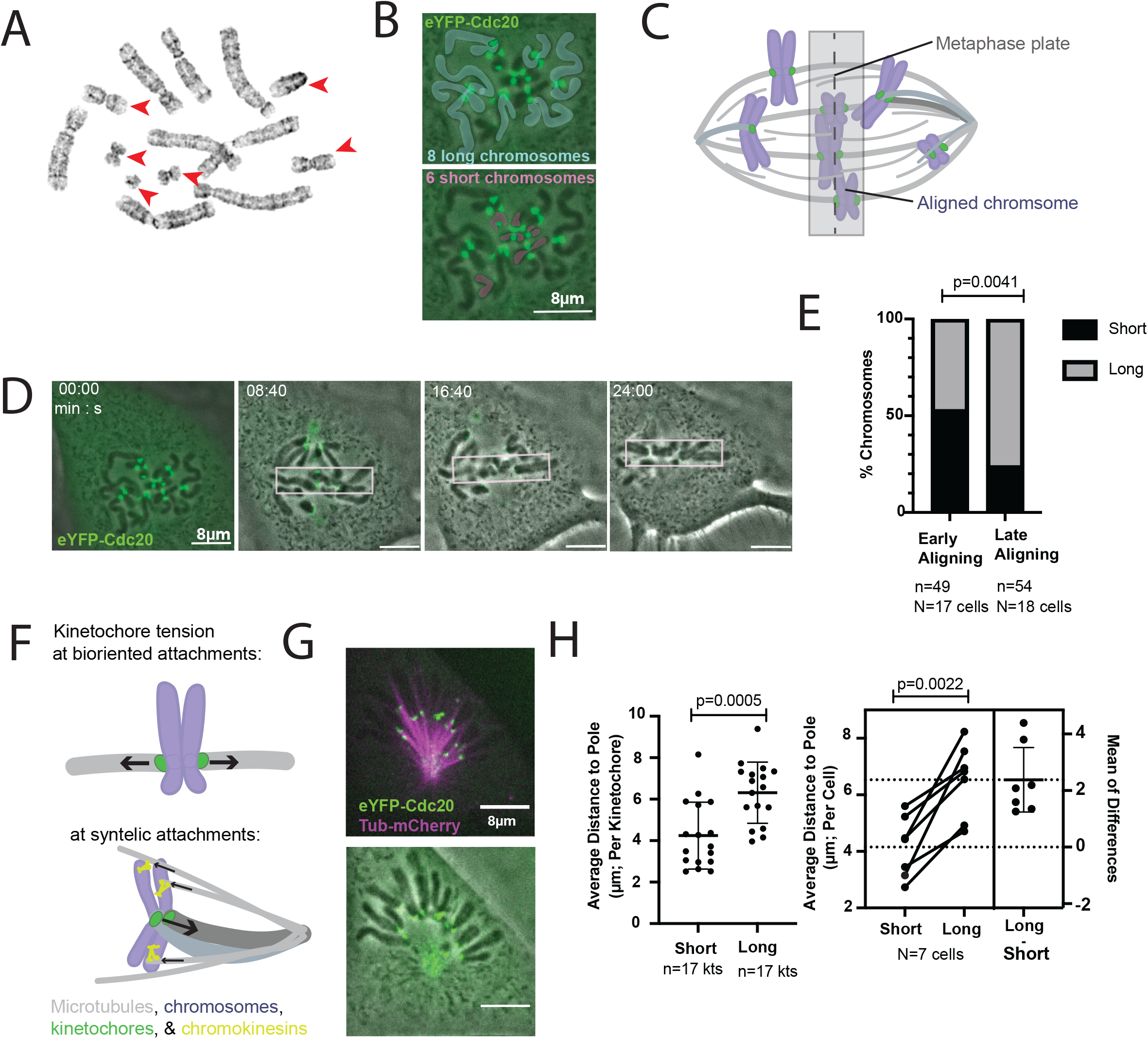
Long chromosomes align less efficiently than short chromosomes and experience higher spindle pushing force. (A) Chromosome spread of PtK2 cell line expressing eYFP-Cdc20. Red arrowheads indicate chromosomes classified as “short”. (B) Long (blue, top) and short (pink, bottom) chromosomes in a live mitotic PtK2 cell. Chromosomes were classified by phase contrast microscopy with the help of temporal tracking information. (C) Spindle assembly schematic depicting aligned, or congressed, chromosomes oscillating within the central gray box of the metaphase plate and unaligned chromosomes, in various aJachment states. Alignment for (D)&(E) is defined by K-K stretch and oscillatory movement within the spindle center as indicated here. (D) Representative time-lapse imaging of spindle assembly of cell shown in (B) showing that some chromosomes align soon after onset of mitosis (pink box indicating oscillatory area of the metaphase plate) while others move to poles, leading to a delay in alignment. See also Video 1. (E) Percent of long and short chromosomes, which are early aligning (the first three to begin oscillating in a given spindle) or late aligning (the last three). n denotes number of chromosomes counted while N denotes number of cells (Fisher’s exact test). (F) For bioriented aJachments, poleward pulling by sister k-fibers produces opposing force that generates kinetochore tension while for syntelic errors, poleward pulling is counteracted by polar ejection force along chromosome arms. To assess whether polar ejection force scales with chromosome size, live imaging was performed on STLC-treated monopolar spindles in PtK2 cells expressing eYFP-Cdc20 and tubulin-mCherry (G) (see also Video 2) and the distance of kinetochores from the pole was used to evaluate the magnitude of pushing force (H) (Unpaired t test).

A longstanding error correction model posits high tension across sister kinetochores as the primary cue that promotes formation of correct attachments while disfavoring incorrect ones (Funabiki, 2019; Sarangapani and Asbury, 2014). Thus, we sought to test whether different sized chromosomes are subject to differing forces. At correct attachments, tension is generated by the opposing force from k-fibers pulling sister kinetochores toward opposite spindle poles; in contrast, incorrect, syntelic attachments (with both sister kinetochores attached to the same spindle pole) may experience kinetochore tension when poleward pulling is counteracted by anti-poleward force along chromosome arms and this may, in turn, affect attachment stability (Fig. 1F). To test how polar ejection forces affect long and short chromosomes, we generated monopolar spindles by treating cells with Eg5 inhibitor STLC and tracked kinetochore movements with respect to the pole. Assuming roughly equal poleward pulling force at all chromosomes due to standard k-fiber size (McEwen et al., 1998, 1997), the distance from poles will be proportional to the strength of the outward polar ejection force. In live monopolar spindles, the kinetochores on long chromosomes were significantly farther from poles on average than short chromosomes (Fig 1G, 1H, and Video 2; unpaired t-test p=0.0005). Thus, long chromosomes biorient less efficiently but also experience higher pushing force than their shorter counterparts.

### Long chromosomes correct errors less efficiently

We hypothesized that increased pushing force at long chromosomes may generate enough tension to stabilize incorrect attachments, prolonging their lifetime and delaying congression. To test this, we enriched for errors in attachment using the Eg5 inhibitor STLC to generate monopolar spindles(Kapoor et al., 2000) and then washed out the drug, allowing spindle bipolarization and error correction to occur (Lampson et al., 2004). Following drug washout, as spindle poles separate, incorrect syntelic attachments (both sister kinetochores attached to the same spindle pole) and incomplete monotelic attachments (one attached sister kinetochore, one unattached) convert to bioriented attachments, which oscillate at the new metaphase plate (Fig. 2A). We measured the time needed for chromosomes to start oscillating after drug washout and used it as a proxy for biorientation and error correction (Fig. 2B & 2C). Alignment times were normalized to measure rank order and account for high variance in time required to achieve bipolarity between different cells (Fig. S1). We find that long chromosomes become bioriented later than short chromosomes following drug washout (Fig. 2C & 2D; Mann-Whitney test p=0.0003). In principle, this could either reflect that long chromosomes have more trouble moving within the spindle than short ones or that long chromosomes correct errors less efficiently. To determine whether drag force limits chromosome movement within the spindle, we measured the velocity of kinetochore movement on long and short chromosomes in monopolar spindles (a prolonged prometaphase-like state) and did not see a significant difference in kinetochore speed between groups (Fig. 2E; p=0.318), consistent with previous findings in grasshopper cells (Nicklas, 1965). Thus, chromosome movement velocity does not impact biorientation. Instead, these data indicate that long chromosomes take longer to correct incorrect syntelic attachments, and together with our previous findings, suggest that this delay might be due to the higher forces they experience.

**Figure 2.**
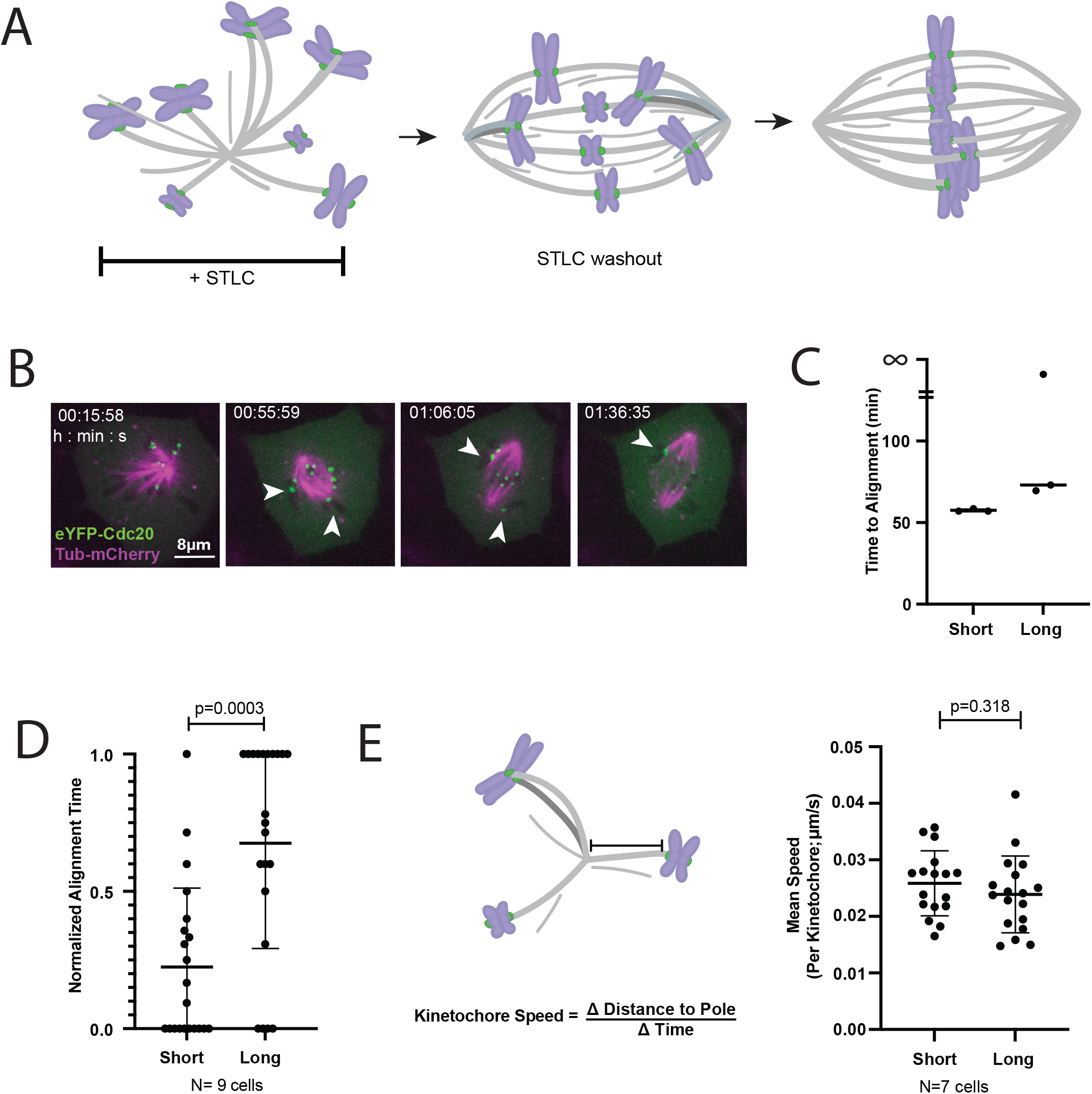
Long chromosomes correct errors less efficiently. (A) Real-time error correction scheme: STLC washout allows monopolar spindles, enriched in aJachment errors, to recover bipolarity and perform error correction. (B) Representative time lapse of error correction assay in PtK2 cells expressing eYFP-Cdc20 and mCherry-α-tubulin. Arrowheads denote long chromosomes stuck in erroneous aJachments. See also Video 3. (C) Raw alignment time of short chromosomes (<7µm) and long chromosomes (≥7µm) in cell shown in (B). (D) Normalized alignment time for population such that time t=0 is the time of the first chromosome oscillating at the metaphase plate following drug washout and t=1 is the time the last chromosome begins oscillating prior to anaphase (Mann-Whitney test). (E) Mean kinetochore speed of long and short chromosomes with respect to the pole in monopolar spindles. (Unpaired t test)

### High polar ejection force increases persistence of polar chromosomes and reduces size-effect

To test if elevated force on long chromosomes is responsible for their delayed error correction, we globally increased polar ejection force and imaged spindle assembly. If elevated force on long chromosomes does not meaningfully contribute to kinetochore tension or attachment stability, increasing anti-poleward force should speed chromosome alignment and reduce dwell time of chromosomes near poles. Indeed, chromokinesins are known to promote chromosome alignment by pushing chromosomes towards the metaphase plate and their depletion slows the progression to metaphase in human cells (Wandke et al., 2012). Alternately, if polar ejection force on long chromosomes can stabilize incorrect attachments, increasing this force globally may result in more delayed chromosome alignment. To discriminate between these possibilities, we perturbed the force balance between poleward pulling at the kinetochore and anti-poleward pushing along chromosome arms by overexpressing the rat kangaroo chromokinesin Kif22/Kid tagged with a HaloTag (Fig. 3A). We confirmed chromokinesin overexpression (Kid OE) and proper localization by immunofluorescence, which showed significantly more total Kid on chromosomes in over expressing cells compared to control (Fig. 3B).

**Figure 3.**
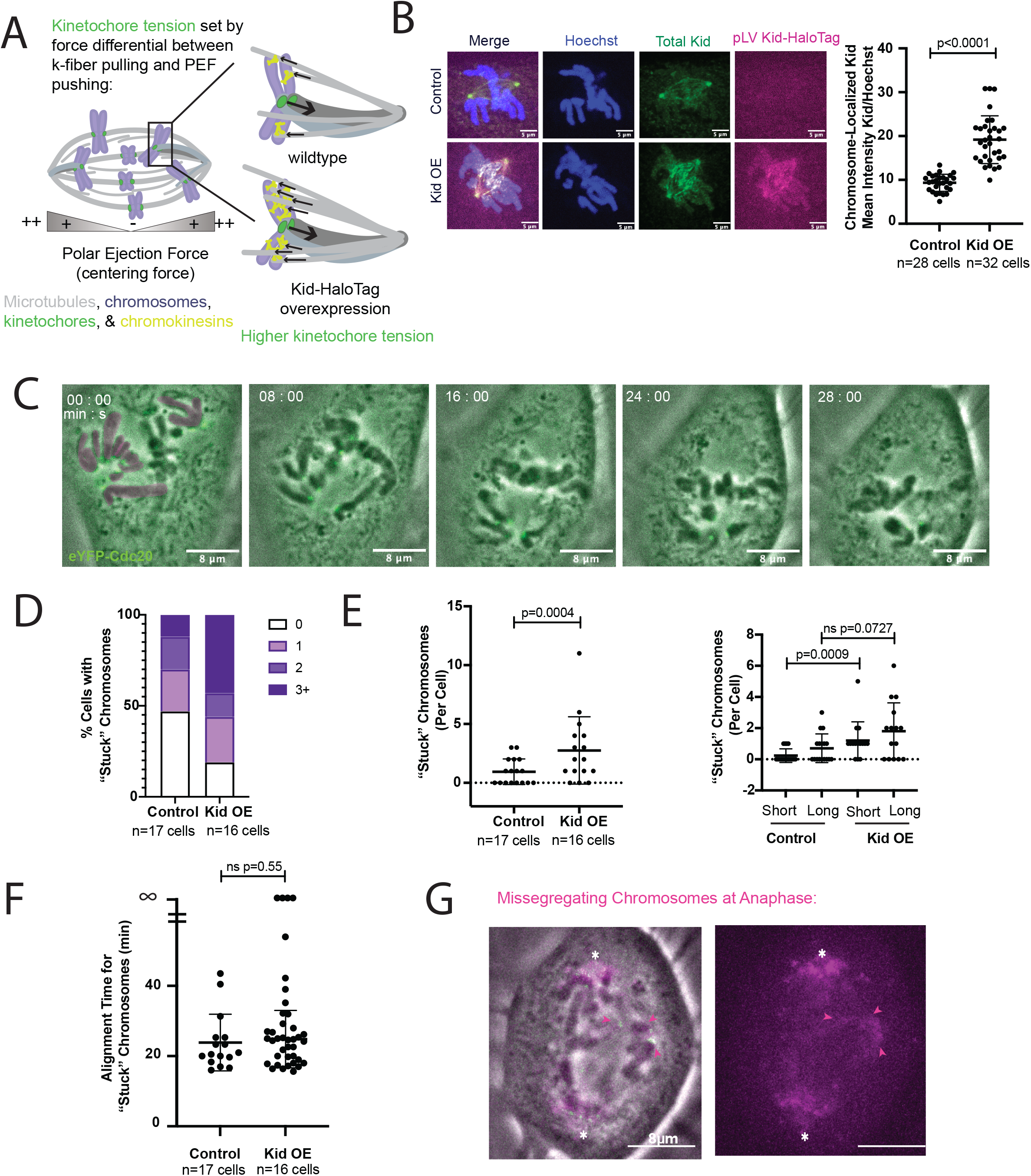
High polar ejection force increases persistence of polar chromosomes and reduces size effect. (A) Polar ejection forces are strongest near poles. Kid overexpression will increase both chromosome centering force exerted by the spindle and, for syntelic kinetochores, kinetochore tension. Tension is set by the force balance between poleward pulling at kinetochores and anti-poleward pushing along chromosome arms. (B) Immunofluorescence of fixed PtK2 cells transiently infected with Kid OE construct (pLV Kid-HaloTag) or not infected and intensity measurements for chromosome-localized Kid normalized to Hoechst signal for one experiment. Experiment was performed in triplicate obtaining similar results each time. (C) Representative time lapse of Kid OE PtK2 cells expressing eYFP-Cdc20. Polar chromosomes are pseudo-colored in magenta for first frame (00:00). See also Video 4. (D) Percent of cells with chromosomes stuck in polar aJachments. Chromosomes were considered stuck if they had not reached alignment within 15 minutes of approaching spindle poles. (E) Number of chromosomes stuck near poles in control and Kid OE spindle assembly (Mann-Whitney test). (F) Time of alignment following pole approach for stuck polar chromosomes in control and Kid OE spindles. (G) Missegregating chromosomes at anaphase in Kid OE spindle from (C). Poles are marked with asterisks and lagging chromosomes denoted by arrowheads.

We imaged spindle assembly and quantified the number of polar chromosomes which persisted at poles for longer than 15 minutes (Fig. 3C and Video 4). Here, we define polar chromosomes as those that did not move to the spindle center but rather oriented sister kinetochores toward the same spindle pole and took on a characteristic V shape of chromosomes under polar ejection force (Fig. 3C). Among control cells, only 53% of cells displayed persistently stuck polar chromosomes compared to 82% of Kid OE cells (Fig. 3D), and on average, Kid OE cells had significantly more chromosomes stuck at poles per cell (Fig. 3E; 0.9 ± 1.1 vs. 2.8 chromosomes ± 2.8, mean±SD; p=0.0004), consistent with observations in Drosophila S2 cells (Cane et al., 2013b). Moreover, when categorized by size, instance of short chromosomes stuck at poles increased significantly with Kid OE (Fig. 3E; 0.2 chromosomes vs. 1.2 chromosomes; p=0.0009), while long chromosomes in persistent polar attachments increased more modestly with no significant difference (0.7 chromosomes vs. 1.7 chromosomes; p=0.07). This distribution suggests a shift in the size-effect delaying chromosome alignment, where rather than a specific delay in long chromosome alignment, a global increase in polar ejection force leads to chromosomes of all sizes getting stuck in polar attachments.

Additionally, consistent with impaired error correction of chromosomes under elevated force, we observed polar chromosome attachments that do not resolve in the duration of imaging (Fig. 3F) and cells entering anaphase with chromosomes stuck at poles as well as lagging chromosomes (Fig. 3G). This indicates the force balance between poleward pulling at the kinetochore and anti-poleward pushing by chromokinesins impacts the ability of cells to detect attachment errors. Despite this, errors that are detected seem to reach alignment on similar timescales to controls (Fig. 3F). Together, we find that globally increasing polar ejection force leads to an enrichment of chromosomes stuck near poles, consistent with a model in which polar ejection force contributes to kinetochore attachment stability even at polar, non-bioriented attachments.

### Chromosome size, not identity, determines chromosome biorientation efficiency

To directly test the role of chromosome size and size-based force in chromosome alignment efficiency, we sought to acutely change chromosome size. To do so, we used laser ablation to cut chromosome arms and measured alignment efficiency. If chromosome size, either directly or indirectly, leads to differential alignment efficiency, shortening chromosome arms should promote earlier chromosome alignment with respect to other long chromosomes in the same cell (model 1; Fig. 4A). If, instead, intrinsic chromosome-specific differences, like kinetochore composition or size (Drpic et al., 2018), dictate chromosome alignment order, ablating chromosome arms should not affect alignment efficiency (model 2; Fig. 4A). We ablated long chromosomes in early prometaphase in cells with many unaligned long chromosomes (Fig. 4B, 4C, and Video 5). Ablated chromosomes congressed to the metaphase plate before other long, unablated chromosomes in the same cell did (Fig. 4D; Mann-Whitney test p=0.004), indicating that chromosome size, not identity, sets alignment order (model 1). This effect was specific to chromosome arm shortening and not due to damaging the arms as control ablations, which nick chromosomes at their ends or along their arms without shortening them (Fig. 4E & 4F), did not change alignment order (Fig. 4G). Given the indistinguishable speeds of kinetochores on long and short chromosomes that exclude any drag effect (Fig. 2E), we conclude that the expedited alignment of ablated chromosomes is the result of reducing polar ejection forces. We propose a model wherein increased polar ejection force on long chromosomes prematurely stabilizes incorrect or incomplete attachments, causing a delay in congression and biorientation (Fig. 5).

**Figure 4.**
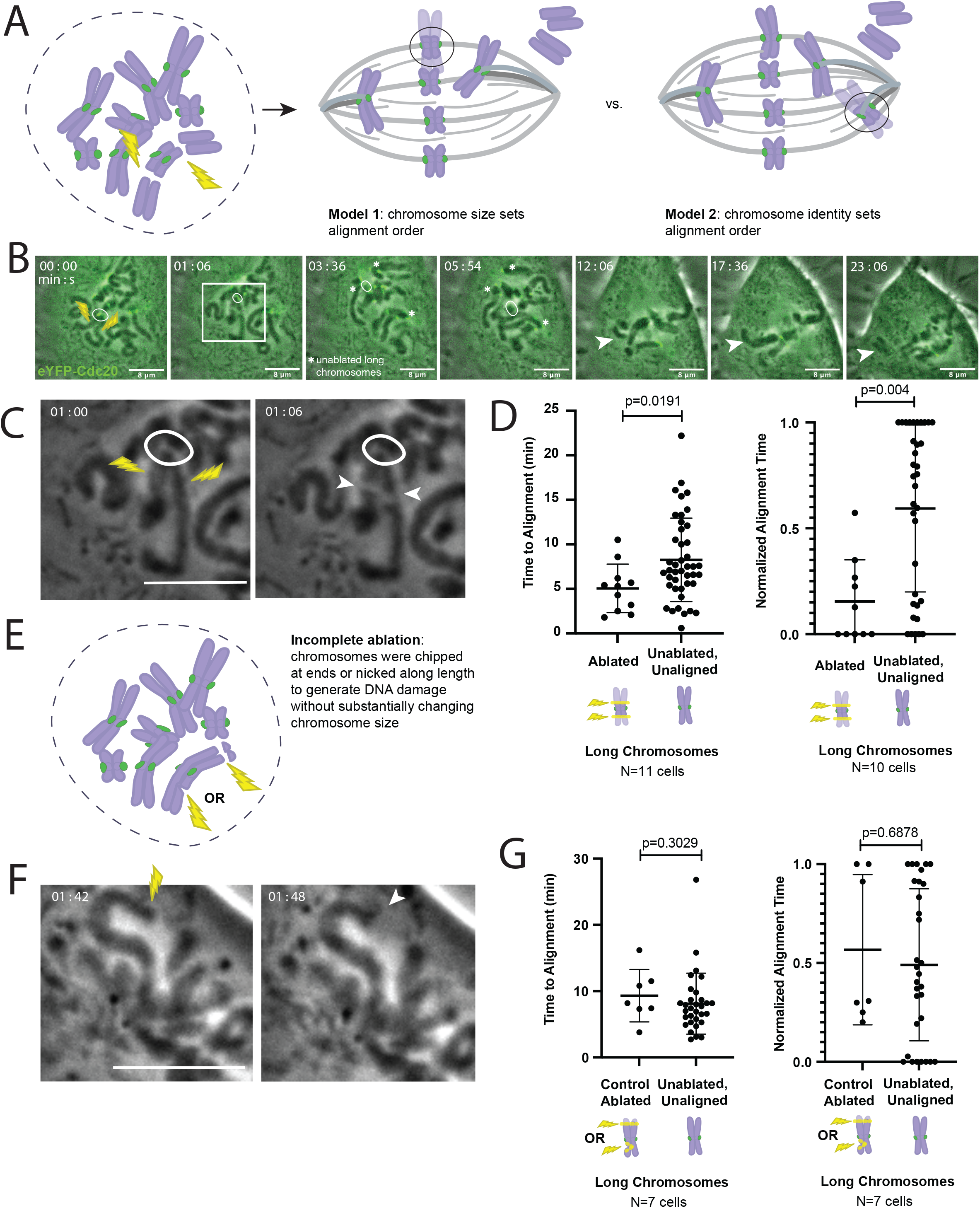
Chromosome size, not identity. determines chromosome biorientation efficiency. (A) Chromosome arm ablation assay: chromosome arms of a long chromosome were cut shortly after nuclear envelope breakdown and kinetochores were followed throughout spindle assembly to determine the time of alignment with respect to other long, unaligned chromosomes in the same cell to test whether chromosome size (model 1; ablated chromosome aligns early) or chromosome identity (model 2; ablated chromosome still aligns late) determine alignment order. (B) Representative example of spindle assembly following chromosome arm ablation in PtK2 cells expressing eYFP-Cdc20. White circle indicates kinetochore on ablated chromosome, arrowheads mark ablated arms, asterisks denote unablated, unaligned long chromosomes. See also Video 5. (C) Zoom in phase contrast images of successful chromosome arm ablation in (B) indicated by white box. Ablation was considered complete if a clear space was observed between the kinetochore-containing fragment and the unaJached arms (arrow heads) and the chromosome fragments were mechanically uncoupled in subsequent frames. (D) Alignment time of ablated and unablated, long chromosomes in raw time and normalized such that t=0 is the first chromosome to align following ablation and t=1 is the last chromosome to align prior to anaphase (Mann-Whitney test). (E) Control ablation scheme depicting how incomplete ablations were performed, either by chipping chromosome ends off or by nicking chromosome arms to damage DNA without changing chromosome size substantially. (F) Representative example of control ablation, chipping chromosome ends. White arrowhead marks chipped chromosome end. (G) Raw time to alignment for control ablations and other unablated long chromosomes in the same cell and normalized alignment time such that t=0 is the first chromosome to align following ablation and t=1 is the last chromosome to align prior to anaphase (Mann-Whitney test).

**Figure 5.**
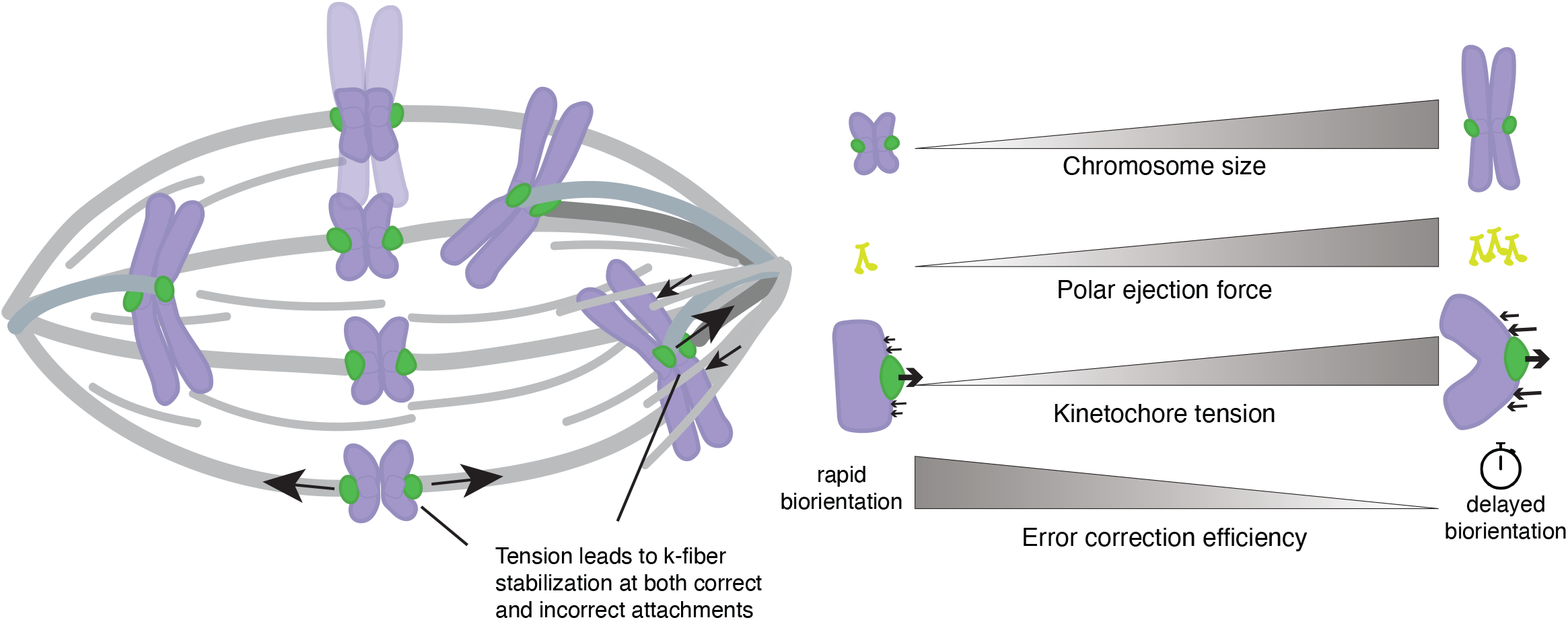
Model for differential tension and error correction efficiency at chromosomes of different sizes. Schematic of an assembling spindle depicting early alignment of short chromosomes and delayed congression of long chromosomes. Based on our findings, we propose that these delays are due to elevated polar ejection force and thus elevated kinetochore tension, which stabilizes both correct and incorrect aJachments. Despite this stabilization, long polar chromosomes typically manage to biorient, suggesting an existing mechanism for detecting errors subject to modest kinetochore tension after some delay. Detection of these errors could occur via a graded tension threshold wherein low tension aJachments are corrected immediately, moderate tension aJachments after a delay, and high tension aJachments after a longer delay. Alternatively, the error correction machinery could rely on a tension-independent mechanism for error detection in these cases.

### Elevated tension at long chromosomes stabilizes incorrect attachments and delays error correction

Accurate chromosome segregation requires cells to be able to distinguish between correct and incorrect attachments, global features of the spindle, despite having only local cues. Here, we ask how the ability both to form bioriented attachments and to detect incorrect attachments varies across chromosomes in the same cell. We demonstrate that in regularly cycling PtK2 cells, with just 14 chromosomes widely ranging in size, the formation of correct, bioriented attachments is biased by chromosome size such that long chromosomes tend to be the last to become correctly attached (Fig. 1). Error correction models have long posited that the kinetochore relies on physical differences—specifically variable tension on kinetochores leading to differential kinetochore phosphorylation—to determine which attachments to reinforce (bioriented) and which to destabilize (syntelic) (Funabiki, 2019; Sarangapani and Asbury, 2014). By enriching for attachment errors, we find that mammalian error correction efficiency varies based on the physical features of the chromosome in question (Fig. 2), and specifically perturbing chromosome size-based force reveals that is the result of the kinetochore sensing and responding to differences in tension (Figs. 3 & 4). Together, these findings suggest that while all chromosomes are competent to form both correct and incorrect attachments, long chromosomes more readily stabilize incorrect attachments, delaying their alignment and biorientation (Fig. 5). In addition to defining biophysical models for error correction, this work defines new questions about protective mechanisms for not just minimizing segregation error rates but also coordinating the threshold for error detection across different chromosomes in the cell.

Overall, this work provides insight into the biophysical mechanism of error correction and reveals that the dynamics of chromosome biorientation are not chromosome-agnostic. Indeed, long chromosomes exhibit distinct behaviors in spindle assembly, more frequently getting stuck at poles (Fig. 3), congressing later in mitosis (Fig. 1), and oscillating with lower amplitude than their short counterparts (Ke et al., 2009). Chromosome arm ablation speeds alignment (Fig. 4D) indicating these are effects of chromosome size and not identity. Still, long and short chromosomes move at similar speeds (Fig. 2E). As such, we propose that differences in alignment order and tendency to dwell at spindle poles stem primarily from chromosome size-based force specifically stabilizing attachments at long polar chromosomes rather than drag force or steric effects. Consistent with this, increasing polar ejection force increases the number of stuck polar chromosomes and reduces the effect of size on tendency to become persistently stuck at poles (Fig. 3D & 3E). Regardless, stuck chromosomes that do align do so on a similar timescale to those in control cells (Fig. 3G), suggesting that beyond a certain tension threshold, additional tension will not slow alignment further but may impact the probability of error detection. This is consistent with work that shows that the tension state impacts kinetochore detachment probability (Chen et al., 2021; De Regt et al., 2022; Miller et al., 2019, 2016; Parmar et al., 2023; Sarangapani et al., 2013). Additionally, attachment stability and error correction efficiency may also be linked to chromosome position in the spindle, leading to differential access to known regulators like Aurora A kinase (Eibes et al., 2023; Ye et al., 2015). Together, these findings support a model in which high force near poles increases the propensity to stabilize polar chromosome attachments (Fig. 5). In healthy cells, elevated polar ejection force at long chromosomes might come from more numerous chromokinesins, non-kinetochore microtubules polymerizing against a larger chromatin surface and generating force, or both (Schneider et al., 2022). Regardless, this size dependent mechanism for error detection is especially interesting considering recent work showing higher rates of missegregation of large chromosomes in human cells (Klaasen et al., 2022; Worrall et al., 2018) and formation of complex attachment errors in polar chromosomes (Vukušić and Tolić, 2022).

Our model, that error correction dynamics depend on chromosome size and resultant kinetochore tension, may have wide ranging functional implications for segregation outcomes, and thus propensity toward aneuploidy. How this differential tension and attachment stability across chromosomes of different sizes alters molecular maturation of the kinetochore in processes such as Ndc80 phosphorylation (Sarangapani et al., 2021), SKA (Cheerambathur et al., 2017; Hanisch et al., 2006) and SKAP (Fang et al., 2009; Schmidt et al., 2010) recruitment, and checkpoint satisfaction (Etemad et al., 2015; Tauchman et al., 2015) remains an exciting open question. It is worth noting that missegregation rates were found to be similar across all chromosomes in PtK1 cells by fixed cell imaging (Torosantucci et al., 2009), yet these results are not inconsistent with a size-based model for error correction efficiency. Chromosome segregation errors rely on failure of both the error correction machinery and the spindle assembly checkpoint (SAC), but how tightly these pathways are coupled may vary across systems. Indeed, observations of SAC satisfaction dynamics have been variable. While Drosophila S2 cells show partial SAC satisfaction at persistently incorrect attachments (Cane et al., 2013a), live monitoring of the SAC in PtK2 cells revealed switch-like loss of Mad1 from the kinetochore (Kuhn and Dumont, 2017). Thus, how and whether persistently stuck polar chromosomes signal the checkpoint in mammalian cells remains unclear. Our work demonstrates that however many errors do or do not slip through at anaphase, the dynamics of error correction do, indeed, vary in PtK2 cells with chromosome size (Fig. 2). Future work will be required to untangle the crosstalk between the SAC and error correction.

Characterizing not just the propensity of individual chromosomes for missegregation but also the features that tune that propensity up or down will inform our understanding of different organisms’ or cell types’ oncogenic capacity and tendency toward aneuploidy. Interesting evolutionary questions arise from non-random missegregation rates. Is there evolutionary pressure to maintain oncogenes on short chromosomes, which may be less likely to missegregate? Or instead, in organisms with long chromosomes carrying oncogenes, is there selective pressure to either homogenize chromosome size over evolutionary time or develop compensatory molecular mechanisms that will equalize missegregation rates? The size-dependence revealed here is likely just one of many features that impact error correction efficiency. Indeed, in different species, chromosome size (Klaasen et al., 2022; Worrall et al., 2018), kinetochore position (metacentric, telocentric, or holocentric) (Dumont et al., 2020), material properties (Cojoc et al., 2016), and size (Drpic et al., 2018), as well as chromatin material properties (Schneider et al., 2022) all have the potential to impact segregation outcomes. Future work on these topics will shed light on how organisms employ either a low threshold for sensing incorrectness, compensatory mechanisms that tune error detection or correction capacity on a per chromosome basis, or both to avoid catastrophic chromosomal instability. Ultimately, the findings herein provide insight into the biophysical mechanism of error detection at the kinetochore and provide a framework through which to think more broadly about how biophysical processes must shape molecular mechanisms across evolution.

## Supplementary Figure Legends

### Video Legends

**Video 1. Long chromosomes align later than short chromosomes during spindle assembly, related to** Fig. 1. Spindle assembly in PtK2 cell expressing eYFP-Cdc20 (kinetochores) to measure chromosome alignment order. Some chromosomes align and begin oscillating soon after the onset of mitosis while others move to poles, delaying their biorientation. Scale bar, 8µm. Video was collected on a spinning disk confocal microscope with fluorescence and phase contrast imaging at one frame every 20 s. Video has been adjusted to play at a constant playback speed of 7 frames/s. Time: h:min:s. Video corresponds to still images from Fig. 1D.

**Video 2. Polar ejection forces push long chromosomes further than short ones from the pole of monopolar spindles, related to** Fig.1. Kinetochore oscillations in a monopolar spindle generated by treating PtK2 cells expressing eYFP-Cdc20 and mCherry-α-tubulin with the Eg5 inhibitor STLC. Average positions of long and short chromosomes were tracked in relation to the pole. Scale bar, 8µm. Video was collected on a spinning disk confocal microscope with fluorescence and phase contrast imaging at one frame every 13 s. Video has been adjusted to play at a constant playback speed of 10 frames/s. Time: h:min:s. Video corresponds to still images from Fig. 1G.

**Video 3. Monopolar spindles regain bipolarity and correct errors following STLC drug washout, related to** Fig. 2. STLC drug washout (time = 0) in PtK2 cell expressing eYFP-Cdc20 and tubulin-mCherry shows spindle recovery from a monopolar to a bipolar state. Error correction was measured by following kinetochore movement and assessing the first timepoint kinetochores begin oscillating at the metaphase plate after washout. Scale bar, 5µm. Video was collected on a spinning disk confocal microscope with fluorescence and phase contrast imaging at one frame every 4 min until 55 min after washout, then every 45 s thereafter. Video has been adjusted to play at a constant playback speed of 7 frames/s. Time: h:min:s. Video corresponds to still images from Fig. 2B.

**Video 4. Chromosomes of all sizes become persistently stuck at poles under high polar ejection force induced by Kid overexpression, related to Fig. 3.** Overexpression of Kid-HaloTag in PtK2 cell expressing eYFP-Cdc20 leads to an increase in chromosomes persistently stuck at poles. Most chromosomes eventually align, and the cell enters anaphase with lagging chromosomes. Scale bar, 8µm. Video was collected on a spinning disk confocal microscope with fluorescentce and phase contrast imaging at one frame every 12s. Video has been adjusted to play at a constant playback speed of 10 frames/s. Time: h:min:s. Video corresponds to still images from Fig. 3C.

**Video 5. Reducing chromosome arm size by laser ablation speeds biorientation during spindle assembly, related to** Fig. 4. Early in mitosis, chromosome arms are severed (time = 1 min, white “X”) from kinetochore-containing fragment (white circle), which proceeds to biorient, oscillating at the metaphase plate prior to other long, unaligned chromosomes (white asterisks). Chromosome arms are mechanically uncoupled from kinetochores as denoted by their presence outside the spindle body at the outset of anaphase (white caret). Laser ablation site marked by white X. Scale bar, 8µm. Video was collected on a spinning disk confocal microscope with fluorescence and phase contrast imaging at one frame every 6s. Video has been adjusted to play at a constant playback speed of 10 frames/s. Time: h:min:s. Video corresponds to still images from Fig. 4B.

## Materials and Methods

### Cell culture

PtK2 cells expressing human eYFP-Cdc20 (gi; from Jagesh Shah, Harvard Medical School, Boston, MA) and wildtype PtK2 cells (ATCC) were cultured at 37°C and 5% CO_2_ in MEM (Invitrogen) supplemented with sodium pyruvate (Invitrogen), non-essential amino acids (Invitrogen), penicillin/streptomycin, and 10% qualified and heat-inactivated fetal bovine serum. Karyotyping and cytogenic analysis were performed on G-banded metaphase cells from the eYFP-Cdc20 PtK2 line by Cell Line Genetics. Cells were transfected using Viafect (Promega) and imaged 72 hours a;er transfection with mCherry-α-tubulin (gi; from M. Davidson, Florida State University, Tallahassee, FL). Transfection reactions were prepared in a 100μl reactions using a 1:6 ratio of DNA to Viafect in OpRMEM media. Overexpression experiments were done with transient lentiviral infection. The coding sequence for the rat kangaroo chromokinesin Kif22/Kid was obtained from the PtK transcriptome (Udy et al., 2015) and cloned into a puromycin resistant lentiviral backbone. Lentivirus was generated in Hek293T cells, and PtK2 cells were infected 48-72 hours prior to imaging spindle assembly.

### Microscopy

Live imaging was done using an inverted (Eclipse Ti-E; Nikon) spinning-disk confocal microscope (CSU-X1; Yokogawa Electric Corporation) with Di01-T405/488/568/647 dichroic head (Semrock), 100x 1.45 Ph3 oil objective, 405 nm (100 mW), 488 nm (120 mW), 561 nm (150 mW), 642 nm (100 mW) diode lasers, emission filters (ET455/50M, ET525/50M, ET630/75M, and ET690/50M; Chroma Technology Corp.), and either an iXon3 camera (Andor Technology) (Fig. 1G, 2B, 3B, 3C, 3G, 4B, 4C, 4E) or a Zyla 4.2 sCMOS camera (Andor Technology) (Fig. 1B, 1D). Cells were plated for imaging on 35-mm dishes with #1.5 poly-D-lysine coated coverslips (MatTek Corporation). For all live experiments except drug washouts, images were collected at bin=1, while drug washout and monopole movies were collected at bin=2 on MetaMorph (7.8, MDS Analytical Technologies) or Micro-Manager (2.0.0). Cells were imaged in phase contrast and fluorescence in a single z-plane (Fig. 3, 4), six z-planes spaced 0.35µm apart (Fig. 1G, 2), or three z-planes spaced 0.35µm apart (Fig. 1B, 1D). Cells were imaged in a humidified stage-top incubation chamber (Tokai Hit) at 37°C with 5% CO_2_. Cells expressing HaloTag constructs were labelled with 200nM JF549 or JF646 (Promega) for 30 minutes prior to imaging.

### Immunofluorescence

To validate chromokinesin overexpression, cells were seeded on poly-L-lysine coated coverslips, transiently infected with lentivirus driving expression of PtK Kid-HaloTag for 48 hours. Cells were then labeled with HaloTag dye JF549 and fixed in 99.8% methanol for 3 minutes at -20°C and permeabilized in TBS1x (Tris buffered saline) + 2% BSA + 0.1% Triton for 30 minutes (IF Buffer herea;er). Primary antibodies were incubated overnight at 4°C at the following concentrations in IF Buffer: mouse anti-α-tubulin (DM1α, 1:1000, Sigma T6199), rabbit anti-rat kangaroo-Kid (3μg/ml, Genscript). The anti-rat kangaroo Kid was raised against the full Kid protein translated from the coding sequence obtained from the PtK transcriptome (Udy et al., 2015). Coverslips were washed three times in IF buffer (5 minutes each) before incubating with secondary antibodies (1:500 in IF buffer, 30 minutes at room temperature): goat anti-rabbit IgG Alexa Fluor 488 (A11008, Invitrogen). Samples were washed three times in IF buffer, incubated with Hoechst 33342 (1μg/ml, H3570, Invitrogen) for one minute, and washed in IF buffer once more prior to mounting on slides with ProLong Gold Antifade Mountant (p36934, Thermo Fisher).

### Laser ablation

Laser ablation experiments were done using 514 nm ns-pulsed laser light and a galvo-controlled MicroPoint Laser System (Andor, Oxford Instruments) operated through Micro-Manager. Spindle assembly was imaged with single z-planes in both phase contrast and 488 nm (to visualize kinetochores expressing eYFP-Cdc20) to identify unaligned long chromosomes (by phase contrast). Chromosomes were selected to ablate if they were relatively isolated (a stretch of chromosome not overlapping with others), not at the spindle center, and if most of the chromosome arms were in focus at a single z-plane (liqle to no tilt). Ablations were performed by firing the laser at 4-8 discrete points across the width of chromosome arms (40-80 pulses of 3ns at 20 Hz) and successful chromosome arm ablations were verified by a visible continuous gap between chromosome segments and mechanical uncoupling of ablated arms and the kinetochore-containing fragment (i.e., moving in different directions). Similarly, control ablations were considered successful if a noticeable change could be seen at the site of laser targeting (a loss of roundness at chromosome ends or visible damage along chromosome arms).

### Drug treatments and washouts

To generate monopolar spindles, Eg5 motor activity was inhibited by addition of S-trityl-L-cysteine (STLC; 7.5μM). Monopoles were imaged and kinetochores tracked for no more than 45 minutes. Error correction was assessed by STLC washout, imaging every four minutes for the first 40-60 minutes until spindles appeared roughly bipolar but had not reached metaphase, then every 20 seconds thereafter.

### Image analysis

Kinetochores were manually tracked using the MTrackJ plugin on FIJI (Schindelin et al., 2012) with a combination of eYFP-Cdc20 and phase contrast to follow individual kinetochores and measure the size of chromosomes. For kinetochores that le; the imaging plane for a short period during a movie, rough position (polar or at the metaphase plate) was recorded using the phase contrast channel. Chromosome size was measured in 2-dimensions using the segmented line tool on FIJI on single z plane phase contrast images to measure the longest dimension of a chromosome. In a crowded spindle, kinetochore tracks were used to find the clearest frame for size measurement. Long chromosomes were considered those ≥7µm and short chromosomes were those <7µm in length.

For alignment order (Fig. 1D) and alignment time (Fig. 2C, 3D, 3E, 4D), the metaphase plate was defined as the region between poles in which oscillating chromosomes resided. Oscillating chromosomes were defined as chromosomes with K-K distance of ≥1.8μm moving periodically toward one spindle pole, then the other. Distance to pole (Fig. 1G) was calculated in monopolar spindles by finding the shortest distance between kinetochore position and pole position (found by MTrackJ using mCherry-α-tubulin fluorescence) and speed of kinetochore movement (Fig. 4E) was determined by measuring the change in kinetochore-to-pole distance over the change in time.

For immunofluorescence, brightness and contrast for each channel were scaled identically within each experiment. A threshold mask (Yen method) was applied using the Hoechst signal (for chromosome selection) to determine the area in which we would measure fluorescence intensity of our protein of interest. Fluorescence intensity was measured of both the chromokinesin Kid and the reference (Hoechst) and normalized by the intensity of the reference (Fig. 3B). Three independent experiments were performed and quantified separately, yielding comparable results.

Statistical analyses were performed in Graphpad Prism 9. For parametric datasets (Fig. 1G), the student’s t test was used. For non-parametric datasets (Fig. 2C, 3B, 3D, 3E, 4D), the Mann-Whitney was used. Fisher’s exact test was used in Fig. 1D.

#### Acknowledgements

We thank Zachary Mullin-Bernstein for help karyotyping. We thank Jagesh Shah for the gi; of the PtK2 eYFP-Cdc20 cell line. We thank Hironori Funabiki, Wallace Marshall, Geeta Narlikar, Orion Weiner, and members of the S. Dumont laboratory for helpful discussions. This work was supported by National Institutes of Health R01GM134132, NIH R35GM136420, National Science Foundation (NSF) Faculty Early Career Development Program 1554139, NSF Center for Cellular Construction DBI-1548297, the Rita Allen Foundation (to S. Dumont), and NIH F31-CA265136-01 and NIH T32EB009383-11A1 (M.K.C.). S. Dumont is a Chan Zuckerberg Biohub investigator.

---

**Figure S1.**
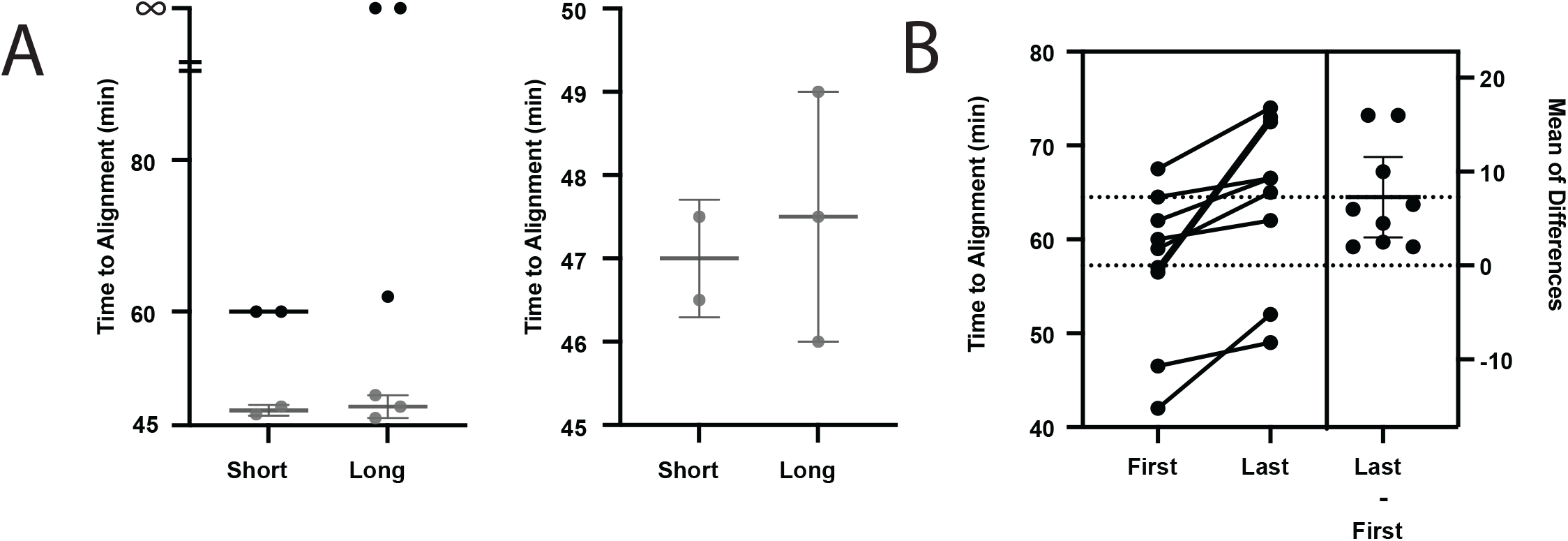
Time to biorientation after STLC washout varies widely due to differences in time to achieve spindle bipolarity. (A) Raw time of chromosome alignment following STLC washout experiments for short and long chromosomes in two independent cells (ex. #1 black and ex. #2 gray; left) and a zoomed in plot for ex. #2 (right) demonstrating that after drug washout, cells took variable amounts of time to achieve spindle bipolarization and chromosome alignment. (B) Time of first detectable oscillating chromosome following STLC washout and last to begin oscillating prior to anaphase within the same cell, showing that some cells had completed chromosome alignment prior to other cells reaching bipolarity. This variable time to become bipolar spindles following STLC washout indicates a need for time normalization in order to compare correction efficiency across cells.

## References

1. Barisic, M., Aguiar, P., Geley, S., Maiato, H., 2014. Kinetochore motors drive congression of peripheral polar chromosomes by overcoming random arm-ejection forces. Nat. Cell Biol. 16, 1249–1256. 10.1038/ncb3060

2. Biggins, S., Murray, A.W., 2001. The budding yeast protein kinase Ipl1/Aurora allows the absence of tension to activate the spindle checkpoint. Genes Dev. 15, 3118–3129. 10.1101/gad.934801

3. Cane, S., McGilvray, P.T., Maresca, T.J., 2013a. Insights from an erroneous kinetochore-microtubule attachment state. BioArchitecture 3, 69–76. 10.4161/bioa.25734

4. Cane, S., Ye, A.A., Luks-Morgan, S.J., Maresca, T.J., 2013b. Elevated polar ejection forces stabilize kinetochore–microtubule attachments. J. Cell Biol. 200, 203–218. 10.1083/jcb.201211119

5. Cheerambathur, D.K., Prevo, B., Hattersley, N., Lewellyn, L., Corbett, K.D., Oegema, K., Desai, A., 2017. Dephosphorylation of the Ndc80 Tail Stabilizes Kinetochore-Microtubule Attachments via the Ska Complex. Dev. Cell 41, 424–437.e4. 10.1016/j.devcel.2017.04.013

6. Cheeseman, I.M., Chappie, J.S., Wilson-Kubalek, E.M., Desai, A., 2006. The Conserved KMN Network Constitutes the Core Microtubule-Binding Site of the Kinetochore. Cell 127, 983–997. 10.1016/j.cell.2006.09.039

7. Chen, G.-Y., Renda, F., Zhang, H., Gokden, A., Wu, D.Z., Chenoweth, D.M., Khodjakov, A., Lampson, M.A., 2021. Tension promotes kinetochore–microtubule release by Aurora B kinase. J. Cell Biol. 220, e202007030. 10.1083/jcb.202007030

8. Cojoc, G., Roscioli, E., Zhang, L., García-Ulloa, A., Shah, J.V., Berns, M.W., Pavin, N., Cimini, D., Tolić, I.M., Gregan, J., 2016. Laser microsurgery reveals conserved viscoelastic behavior of the kinetochore. J. Cell Biol. 212, 767–776. 10.1083/jcb.201506011

9. De Regt, A.K., Clark, C.J., Asbury, C.L., Biggins, S., 2022. Tension can directly suppress Aurora B kinase-triggered release of kinetochore-microtubule attachments. Nat. Commun. 13, 2152. 10.1038/s41467-022-29542-8

10. DeLuca, J.G., Gall, W.E., Ciferri, C., Cimini, D., Musacchio, A., Salmon, E.D., 2006. Kinetochore Microtubule Dynamics and Attachment Stability Are Regulated by Hec1. Cell 127, 969– 982. 10.1016/j.cell.2006.09.047

11. DeLuca, K.F., Lens, S.M.A., DeLuca, J.G., 2011. Temporal changes in Hec1 phosphorylation control kinetochore–microtubule attachment stability during mitosis. J. Cell Sci. 124, 622–634. 10.1242/jcs.072629

12. Drpic, D., Almeida, A.C., Aguiar, P., Renda, F., Damas, J., Lewin, H.A., Larkin, D.M., Khodjakov, A., Maiato, H., 2018. Chromosome Segregation Is Biased by Kinetochore Size. Curr. Biol. 28, 1344–1356.e5. 10.1016/j.cub.2018.03.023

13. Dumont, M., Gamba, R., Gestraud, P., Klaasen, S., Worrall, J.T., De Vries, S.G., Boudreau, V., Salinas-Luypaert, C., Maddox, P.S., Lens, S.M., Kops, G.J., McClelland, S.E., Miga, K.H., Fachinetti, D., 2020. Human chromosome-specific aneuploidy is influenced by DNA - dependent centromeric features. EMBO J. 39, e102924. 10.15252/embj.2019102924

14. Eibes, S., Rajendraprasad, G., Guasch-Boldu, C., Kubat, M., Steblyanko, Y., Barisic, M., 2023. CENP-E activation by Aurora A and B controls kinetochore fibrous corona disassembly. Nat. Commun. 14, 5317. 10.1038/s41467-023-41091-2

15. Etemad, B., Kuijt, T.E.F., Kops, G.J.P.L., 2015. Kinetochore–microtubule attachment is sufficient to satisfy the human spindle assembly checkpoint. Nat. Commun. 6, 8987. 10.1038/ncomms9987

16. Fang, L., Seki, A., Fang, G., 2009. SKAP associates with kinetochores and promotes the metaphase-to-anaphase transition. Cell Cycle 8, 2819–2827. 10.4161/cc.8.17.9514

17. Funabiki, H., 2019. Correcting aberrant kinetochore microtubule attachments: a hidden regulation of Aurora B on microtubules. Curr. Opin. Cell Biol. 58, 34–41. 10.1016/j.ceb.2018.12.007

18. Hanisch, A., Silljé, H.H.W., Nigg, E.A., 2006. Timely anaphase onset requires a novel spindle and kinetochore complex comprising Ska1 and Ska2. EMBO J. 25, 5504–5515. 10.1038/sj.emboj.7601426

19. Iemura, K., Tanaka, K., 2015. Chromokinesin Kid and kinetochore kinesin CENP-E differentially support chromosome congression without end-on attachment to microtubules. Nat. Commun. 6, 6447. 10.1038/ncomms7447

20. Kapoor, T.M., Mayer, T.U., Coughlin, M.L., Mitchison, T.J., 2000. Probing Spindle Assembly Mechanisms with Monastrol, a Small Molecule Inhibitor of the Mitotic Kinesin, Eg5. J. Cell Biol. 150, 975–988. 10.1083/jcb.150.5.975

21. Ke, K., Cheng, J., Hunt, A.J., 2009. The Distribution of Polar Ejection Forces Determines the Amplitude of Chromosome Directional Instability. Curr. Biol. 19, 807–815. 10.1016/j.cub.2009.04.036

22. Klaasen, S.J., Truong, M.A., van Jaarsveld, R.H., Koprivec, I., Štimac, V., de Vries, S.G., Risteski, P., Kodba, S., Vukušić, K., de Luca, K.L., Marques, J.F., Gerrits, E.M., Bakker, B., Foijer, F., Kind, J., Tolić, I.M., Lens, S.M.A., Kops, G.J.P.L., 2022. Nuclear chromosome locations dictate segregation error frequencies. Nature 607, 604–609. 10.1038/s41586-022-04938-0

23. Kuhn, J., Dumont, S., 2017. Spindle assembly checkpoint satisfaction occurs via end-on but not lateral attachments under tension. J. Cell Biol. 216, 1533–1542. 10.1083/jcb.201611104

24. Lampson, M., Grishchuk, E., 2017. Mechanisms to Avoid and Correct Erroneous Kinetochore-Microtubule Attachments. Biology 6, 1. 10.3390/biology6010001

25. Lampson, M.A., Renduchitala, K., Khodjakov, A., Kapoor, T.M., 2004. Correcting improper chromosome–spindle attachments during cell division. Nat. Cell Biol. 6, 232–237. 10.1038/ncb1102

26. Levesque, A.A., Compton, D.A., 2001. The chromokinesin Kid is necessary for chromosome arm orientation and oscillation, but not congression, on mitotic spindles. J. Cell Biol. 154, 1135–1146. 10.1083/jcb.200106093

27. McEwen, B.F., Ding, Y., Heagle, A.B., 1998. Relevance of kinetochore size and microtubule-binding capacity for stable chromosome attachment during mitosis in PtK1 cells. Chromosome Res. 6, 123–132. https://doi-org.ucsf.idm.oclc.org/10.1023/A:1009239013215

28. McEwen, B.F., Heagle, A.B., Cassels, G.O., Buttle, K.F., Rieder, C.L., 1997. Kinetochore Fiber Maturation in PtK1 Cells and Its Implications for the Mechanisms of Chromosome Congression and Anaphase Onset. J. Cell Biol. 137, 1567–1580. 10.1083/jcb.137.7.1567

29. Miller, M.P., Asbury, C.L., Biggins, S., 2016. A TOG Protein Confers Tension Sensitivity to Kinetochore-Microtubule Attachments. Cell 165, 1428–1439. 10.1016/j.cell.2016.04.030

30. Miller, M.P., Evans, R.K., Zelter, A., Geyer, E.A., MacCoss, M.J., Rice, L.M., Davis, T.N., Asbury, C.L., Biggins, S., 2019. Kinetochore-associated Stu2 promotes chromosome biorientation in vivo. PLOS Genet. 15, e1008423. 10.1371/journal.pgen.1008423

31. Musacchio, A., Desai, A., 2017. A Molecular View of Kinetochore Assembly and Function. Biology 6, 5. 10.3390/biology6010005

32. Nicklas, R.B., 1965. CHROMOSOME VELOCITY DURING MITOSIS AS A FUNCTION OF CHROMOSOME SIZE AND POSITION. J. Cell Biol. 25, 119–135. 10.1083/jcb.25.1.119

33. Nicklas, R.B., Koch, C.A., 1969. CHROMOSOME MICROMANIPULATION. J. Cell Biol. 43, 40–50. 10.1083/jcb.43.1.40

34. Parmar, S., Gonzalez, S.J., Heckel, J.M., Mukherjee, S., McClellan, M., Clarke, D.J., Johansson, M., Tank, D., Geisness, A., Wood, D.K., Gardner, M.K., 2023. Robust microtubule dynamics facilitate low-tension kinetochore detachment in metaphase. J. Cell Biol. 222, e202202085. 10.1083/jcb.202202085

35. Powers, A.F., Franck, A.D., Gestaut, D.R., Cooper, J., Gracyzk, B., Wei, R.R., Wordeman, L., Davis, T.N., Asbury, C.L., 2009. The Ndc80 Kinetochore Complex Forms Load-Bearing Attachments to Dynamic Microtubule Tips via Biased Diffusion. Cell 136, 865–875. 10.1016/j.cell.2008.12.045

36. Rieder, C., Salmon, E., 1994. Motile kinetochores and polar ejection forces dictate chromosome position on the vertebrate mitotic spindle. J. Cell Biol. 124, 223–233. 10.1083/jcb.124.3.223

37. Rieder, C.L., Davison, E.A., Jensen, L.C., Cassimeris, L., Salmon, E.D., 1986. Oscillatory movements of monooriented chromosomes and their position relative to the spindle pole result from the ejection properties of the aster and half-spindle. J. Cell Biol. 103, 581–591. 10.1083/jcb.103.2.581

38. Sarangapani, K.K., Akiyoshi, B., Duggan, N.M., Biggins, S., Asbury, C.L., 2013. Phosphoregulation promotes release of kinetochores from dynamic microtubules via multiple mechanisms. Proc. Natl. Acad. Sci. 110, 7282–7287. 10.1073/pnas.1220700110

39. Sarangapani, K.K., Asbury, C.L., 2014. Catch and release: how do kinetochores hook the right microtubules during mitosis? Trends Genet. 30, 150–159. 10.1016/j.tig.2014.02.004

40. Sarangapani, K.K., Koch, L.B., Nelson, C.R., Asbury, C.L., Biggins, S., 2021. Kinetochore-bound Mps1 regulates kinetochore–microtubule attachments via Ndc80 phosphorylation. J. Cell Biol. 220, e202106130. 10.1083/jcb.202106130

41. Schmidt, J.C., Kiyomitsu, T., Hori, T., Backer, C.B., Fukagawa, T., Cheeseman, I.M., 2010. Aurora B kinase controls the targeting of the Astrin–SKAP complex to bioriented kinetochores. J. Cell Biol. 191, 269–280. 10.1083/jcb.201006129

42. Schneider, M.W.G., Gibson, B.A., Otsuka, S., Spicer, M.F.D., Petrovic, M., Blaukopf, C., Langer, C.C.H., Batty, P., Nagaraju, T., Doolittle, L.K., Rosen, M.K., Gerlich, D.W., 2022. A mitotic chromatin phase transition prevents perforation by microtubules. Nature 609, 183–190. 10.1038/s41586-022-05027-y

43. Tauchman, E.C., Boehm, F.J., DeLuca, J.G., 2015. Stable kinetochore–microtubule attachment is sufficient to silence the spindle assembly checkpoint in human cells. Nat. Commun. 6, 10036. 10.1038/ncomms10036

44. Thompson, A.F., Blackburn, P.R., Arons, N.S., Stevens, S.N., Babovic-Vuksanovic, D., Lian, J.B., Klee, E.W., Stumpff, J., 2022. Pathogenic mutations in the chromokinesin KIF22 disrupt anaphase chromosome segregation. eLife 11, e78653. 10.7554/eLife.78653

45. Torosantucci, L., De Santis Puzzonia, M., Cenciarelli, C., Rens, W., Degrassi, F., 2009. Aneuploidy in mitosis of PtK1 cells is generated by random loss and nondisjunction of individual chromosomes. J. Cell Sci. 122, 3455–3461. 10.1242/jcs.047944

46. Vukušić, K., Tolić, I.M., 2022. Polar Chromosomes—Challenges of a Risky Path. Cells 11, 1531. 10.3390/cells11091531

47. Wan, X., Cimini, D., Cameron, L.A., Salmon, E.D., 2012. The coupling between sister kinetochore directional instability and oscillations in centromere stretch in metaphase PtK1 cells. Mol. Biol. Cell 23, 1035–1046. 10.1091/mbc.e11-09-0767

48. Wandke, C., Barisic, M., Sigl, R., Rauch, V., Wolf, F., Amaro, A.C., Tan, C.H., Pereira, A.J., Kutay, U., Maiato, H., Meraldi, P., Geley, S., 2012. Human chromokinesins promote chromosome congression and spindle microtubule dynamics during mitosis. J. Cell Biol. 198, 847–863. 10.1083/jcb.201110060

49. Worrall, J.T., Tamura, N., Mazzagatti, A., Shaikh, N., van Lingen, T., Bakker, B., Spierings, D.C.J., Vladimirou, E., Foijer, F., McClelland, S.E., 2018. Non-random Mis-segregation of Human Chromosomes. Cell Rep. 23, 3366–3380. 10.1016/j.celrep.2018.05.047

50. Ye, A.A., Deretic, J., Hoel, C.M., Hinman, A.W., Cimini, D., Welburn, J.P., Maresca, T.J., 2015. Aurora A Kinase Contributes to a Pole-Based Error Correction Pathway. Curr. Biol. 25, 1842–1851. 10.1016/j.cub.2015.06.021

51. Zaytsev, A.V., Mick, J.E., Maslennikov, E., Nikashin, B., DeLuca, J.G., Grishchuk, E.L., 2015. Multisite phosphorylation of the NDC80 complex gradually tunes its microtubule-binding affinity. Mol. Biol. Cell 26, 1829–1844. 10.1091/mbc.E14-11- 1539

